# Super-resolution segmentation network for reconstruction of packed neurites

**DOI:** 10.1101/2020.06.09.143347

**Authors:** Zhou Hang, Quan Tingwei, Huang Qing, Liu Tian, Cao Tingting, Zeng Shaoqun

## Abstract

Neuron reconstruction can provide the quantitative data required for measuring the neuronal morphology and is crucial in the field of brain research. However, the difficulty in reconstructing packed neuritis, wherein massive labor is required for accurate reconstruction in most cases, has not been resolved. In this work, we provide a fundamental pathway for solving this challenge by proposing the use of the super-resolution segmentation network (SRSNet) that builds the mapping of the neurites in the original neuronal images and their segmentation in a higher-resolution space. SRSNet focuses on enlarging the distances between the boundaries of the packed neurites producing the high-resolution segmentation images. Thus, in the construction of the training datasets, only the traced skeletons of neurites are required, which vastly increase the usability of SRSNet. From the results of the experiments conducted in this work, it has been observed that SRSNet achieves accurate reconstruction of packed neurites where the other state-of-the-art methods fail.

## Introduction

A neuron cell is the basic structural unit in neuronal circuits [1]. Observing and measuring the neuronal morphology plays a vital role in the field of brain research [2]. A series of advances in the optical imaging and molecular labeling techniques have enabled us to observe an almost complete neuronal morphology with sub-cellular resolution and have pushed the understanding of neuronal anatomy to a new level [3–5]. However, improvements to the measurements of the observed structure can still be done. This is because the difficulties that occur in reconstructing dense neurites, which is an indispensable ingredient in the measurement of neuronal morphology, has not been resolved satisfactorily, and many reconstruction errors require manual revision.

The procedure of reconstructing dense neurites is an open question in the subject of neuronal image analysis. Its challenge originates from the following facts: A neurite and its nearest neighbors have a small distance between their boundaries and their structure thus creates confusion when using the currently available reconstruction methods. The crossover neurites form a variety of potential link patterns whose number increases exponentially with the number of neurites. This makes it difficult to identify the correct links in the crossover region. Furthermore, the effect of the point spread function, the variations in the image intensity and radius along the length of each neurite, and the inhomogeneous background worsen the reconstruction accuracy of dense neurites.

Several methods have been proposed for this reconstruction process. The common characteristic among these methods is their focus on identifying the false links between the traced neurites. Graph partition or machine learning based methods are generally employed to search for the optimal links between the traced neurites which makes the morphological features of the reconstructed neurites to be consistent with certain summarized rules [6–11]. However, the dense distribution of neurites and a low image quality usually result in numerous and complicated false links. The real links are difficult to extract and find using the algorithms. Even if the different scales of the morphological features and the positions of somas are fully considered, a substantial part of the neurites cannot be correctly assigned to their own neurons, especially for ones deviating from somas. The fundamental pathway to resolving this problem of reconstructing dense neurites is to improve the neuronal image resolution to be able to separate the densely packed neurites. Developing the optical system or super-resolution image reconstruction methods in principle is feasible, but these methods are expensive.

In this work, we propose a super-resolution segmentation network (SRSNet) for the reconstruction of dense neurites. This network inherits the advantages of the super-resolution image reconstruction network, enlarges the distances between the boundaries of the neurites and directly outputs the segmentation of the dense neurites in a high-resolution (HR) image space. The proposed network does not require massive three-dimensional (3D) registrations of high-resolution images, which are necessary for super-resolution image reconstruction network, but which are difficult to obtain in practice. Compared to the original image stack, the total voxel of its super-resolution segmentation image stack has a 16-fold improvement, and approximately 50 % decrease is achieved in the radius of the neurites. We demonstrate the vast enhancement that SRSNet provides in the reconstruction accuracy of dense neurites. For some testing datasets that the current methods fail to reconstruct, the reconstruction precision achieved by employing SRSNet ranges from 0.77 to 0.99. Overall, SRSNet essentially reduces the complexity of neuronal images and provides a completely new pathway for solving the problem of reconstructing dense neurites.

## Methods

Image resolution is a key factor in determining the reconstruction accuracy of densely distributed neurites. Limited by the imaging system in practice, the image resolution is not sufficiently high to separate the densely distributed neurites in the observed image stacks. This also hinders the improvement of image resolution with deep convolution networks, which require massive HR images for constructing the training datasets. SRSNet breaks the rule that an HR imaging system is necessary for dense neurite reconstruction, and transforms the neuronal images into a segmentation image in a higher-resolution space in which the densely distributed neurites are separated. To illustrate this point, we introduce the SRSNet structure, the training datasets construction, and some related materials.

In general, for an optical imaging system, the observed image can be approached by convoluting an HR image with the point spread function (PSF) and then downsampling the convoluted image. The PSF is determined by the optical system and keeps stable. Thus, the different radii of neurites can be identified in the HR image, which are, in general, difficult to estimate in practice. In the design of SRSNet, we assume that the convolution of the neurites with a fixed radius and a population of PSFs also approach the observed image (Fig. 1(a)). This assumption is based on the basic property of the convolution operation and makes it easy to generate an HR image corresponding to its observed image. The generated HR image consists of a series of cylinders that connect with each other and the center-point curves of which are the traced skeletons of neurites in the observed image. In this way, we build the mapping of the observed image and its corresponding HR image.

**Fig. 1.**
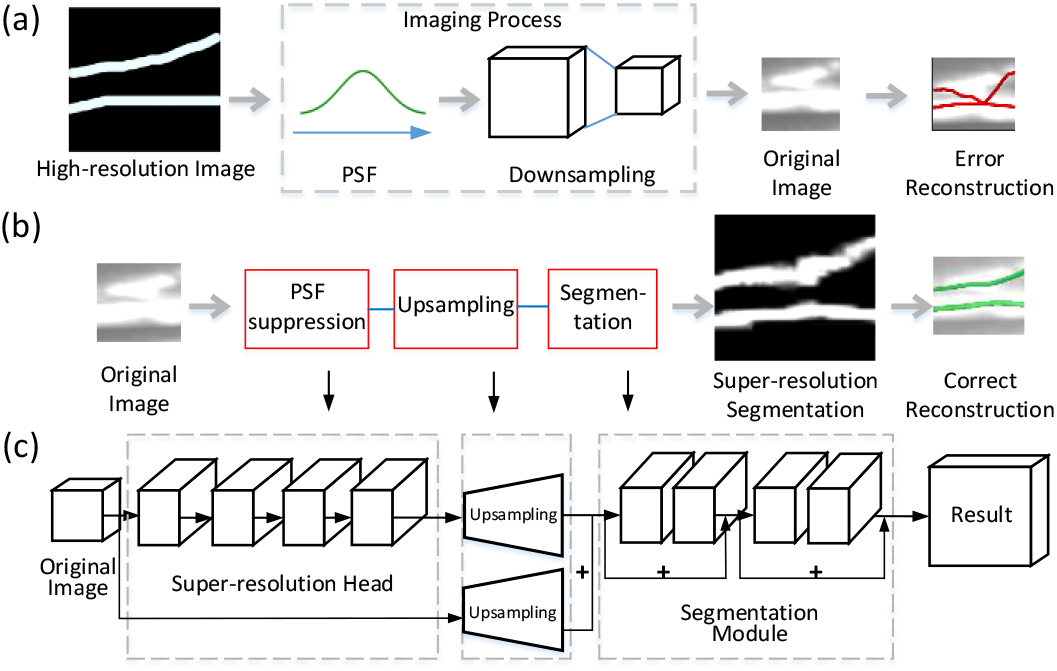
Block diagram showing the structure of SRSNet. (**a**) The imaging model describes the mapping of an HR image and its observed image, which supports the design of the SRSNet. For the observed image stacks, its low resolution results in the error in neurite reconstruction. (**b**) The transformation from the observed image stack to its HR segmentation image stack obtained by SRSNet in which the two densely distributed neurites are separated. (**c**) The SRSNet includes three subnetworks. The two former subnetworks are used for extracting HR feature image stacks. The final subnetwork is used for segmenting the HR feature image stacks.

For capturing the specific mapping as described above, SRSNet is designed to have a cascade of super-resolution head, upsampling, and segmentation networks (**Figs. 1(b) and 1(c)**). The super-resolution head network [12] is used to extract the feature map of the observed image and its corresponding HR image. Its structure is a small residual neural network (ResNet) [13] and consists of one convolution layer and four linear low-rank layers [12]. The upsampling network consists of two upsampling layers. One layer is the pixel shuffle 3D [14] used for increasing the voxels of the feature image stack generated by the super-resolution head network. As the resolution of z-axis of real neuronal images is higher than that of x- and y-axis, all the feature image stacks are enlarged by 2x, 2x, and 4x along the x-, y-, and z-axis respectively. The other layer performs an operation of upsampling the original image stack using the trilinear method and convolutes the upsampled image with 128 template kernels, which indicates 128 feature image stacks. All the feature image stacks generated using these two upsampling networks are summed as the input to the segmentation network which ensures that the final output image stacks coincide with the training set in which the HR-image stack generated with its corresponding observed image stack is binary. Its structure is common and contains four ResNet layers, a convolution layer, and a sigmoid layer.

The training dataset is generated as follows: In a given image stack, the neurites are traced and the traced skeletons are optimized for further approaching the center-lines of the neurites. The positions of these skeletons are mapped in the HR image stack which is generated by dividing each voxel equally in the original image into 2 × 2 × 4 voxels. In the HR image stack, the traced skeletons are resampled to make the distance between any two neighboring skeleton points equal to one voxel size and the 26 neighboring regions of each resampled skeleton point are labeled. The labeled region is regarded as the segmentation of neurites in the HR image stack. Using the above procedure, we obtained sample pairs consisting of a neuronal image stack and its HR segmentation image stack. We used 87 image stacks which included crossover neurites and their HR segmentation image stacks to construct the training datasets. Each image stack and its HR segmentation image stack have 48 × 48 × 24 and 96 × 96 × 96 voxels respectively.

In the training phase, we randomly clipped the image stack with 24 × 24 × 16 voxels and its corresponding HR segmentation image stack with 48 × 48 × 64 from the training datasets and then flipped the image pairs along the x, y or z direction. These argumentation images were used for training the SRSNet. The Adam optimizer [15] was used with a learning rate of 2 × 10^−4^. Assuming that the input image stack with 24 × 24 × 16 voxels occupies 4.3 GB video memory in the training phase, the batch size was set to 1. The training required approximately 17 h on a computing platform having Windows 10 operating system, Intel CPU i5 8300H, 8 GB memory, and GTX 1060 (6 GB). SRSNet was constructed using Pytorch.

In the prediction phase, we divided an image stack into sub-image stacks, all with 32 × 32 × 32 voxels. It occupied 3.3 GB video memory. In this work, any two neighboring sub-image stacks had an overlap region of 16 voxels. To acquire the best segmentation, the image part that was 8 voxels away from boundary was replaced by the corresponding image part in the neighboring sub-image stacks. The central part of the predicted image, which was 1/8th of the total volume, was taken to be valid. The overlap region could be smaller in other works. After prediction, these sub HR segmentation parts were merged into a whole HR segmentation image block and the overlap region was processed using non-maximum suppression. The prediction of a 128 × 128 × 64 voxels image stack requires approximately 380 s, i.e., 2.2 × 10^4^ voxels per second, considering the overlap region. The computing platform was the same as in the training phase, i.e., GTX 1060 (6 GB).

The total loss function of SRSNet consists of focal loss and soft dice loss. The soft dice loss is defined as

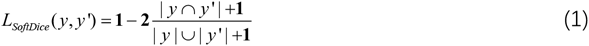

where y denotes the predicted super-resolution segmentation and y′ denotes the segmentation ground truth. Both the numerator and denominator are added with 1 to avoid the all-zero exception. The focal loss is defined as

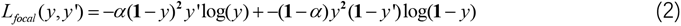

Similar to the study by Lin, et al. [16] we have set the power of the predicted segmentation item to the recommended value of 2 as this will enhance the learning of hard samples. The parameter *α* has been used to balance the positive and negative samples. In the training phase, an important key point was to set *α* in the interval [0.8,0.95]. The value of *α* could be decreased to 0.7 gradually. If its initial value was lower than 0.5, it would result in poor segmentation results. The total loss is defined as

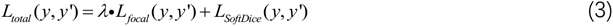

Because of the small portion of positive signals in the neuronal images, the value of the focal loss was small. In contrast, the proportion of positive and negative sample had little influence on the soft dice loss. We have introduced the parameter *λ* for balancing the two loss functions in the total loss. In this work, the value of *λ* has been set to 50.

We have introduced a new neurite tracing evaluation index for evaluating the accuracy of the neurite skeleton extraction and the relationship of the neurite connection after tracing. On the basis of the connection relationship of the neurites, we clustered the connected neurites into a single cluster, called the neurite cluster. One neuronal image contained several neurite clusters. For the single neurite cluster ground truth, we matched each of the neurite clusters, which were traced by automatic tracing methods. The evaluation metrics, namely, precision and recall, were calculated by comparing the best matched neurite cluster skeleton and the neurite cluster ground truth. The length of the neurite clusters was calculated and normalized. We multiplied the values of precision and recall by the normalized length and summed them to obtain the total value of precision and recall. The total precision and recall of all neurite clusters in a neuronal image can evaluate the processes of neurite skeleton extraction and neurite connection relationship simultaneously.

## Results

We have carried on three experiments to evaluate the segmentation performance of SRSNet. In the first experiment, we applied SRSNet on synthetic data to produce super-resolution segmentation and compared the tracing accuracy of previous automatic tracing methods on synthetic data which have dense network of neurites. In the last two experiment, we obtained the qualitative tracing analysis of super-resolution segmentation from mouse neuronal images acquired by fMOST [3].

Second, we demonstrate the tracing performance of super-resolution segmentation on the synthesized chaotic images, which have dense neurites network (**Fig. 2**). The size of the simulation image is 256 × 256 × 128 voxels and its resolution ratio is set as 0.2 × 0.2 × 1 μm. The size of the super-resolution segmentation is 512 × 512 × 512 voxels. We have synthesized five groups of chaotic neurite images. There are three images in each groups. The simulation images have been sorted according to the density of neurites. The minimum and maximum average densities of the five groups are 3.44 m/mm^3^ and 54.61 m/mm^3^ respectively. The simulation image in **Fig. 2(a)** refer to image group 3 in **Fig. 2(b)**. The density of image group 3 is equal to the maximum density of real neuronal images in the following experiment. It means that the simulation image in **Fig. 2(a)** can be regarded as the densest image from our real neuronal dataset. The corresponding traced super-resolution image and the tracing result obtained have also been shown in the figure. The tracing precision of the super-resolution segmentation in **Fig. 2(a)** is 0.82, while its tracing recall is 0.94. The other four tracing methods have been applied on the original images. The tracing recall values of Neutube [17], Snake [18], PHDF [19], and NeuroGPS-Tree [8] methods are 0.86, 0.90, 0.99, and 0.97 respectively. In **Fig. 2(b)**, the tracing precision of super-resolution segmentation images is always higher than four other methods, and achieve precision value of 0.56 in group fifth, while that of four other methods is about 0.1. These tracing methods connect most of the neurites into one cluster, resulting in poor precision.

**Fig. 2.**
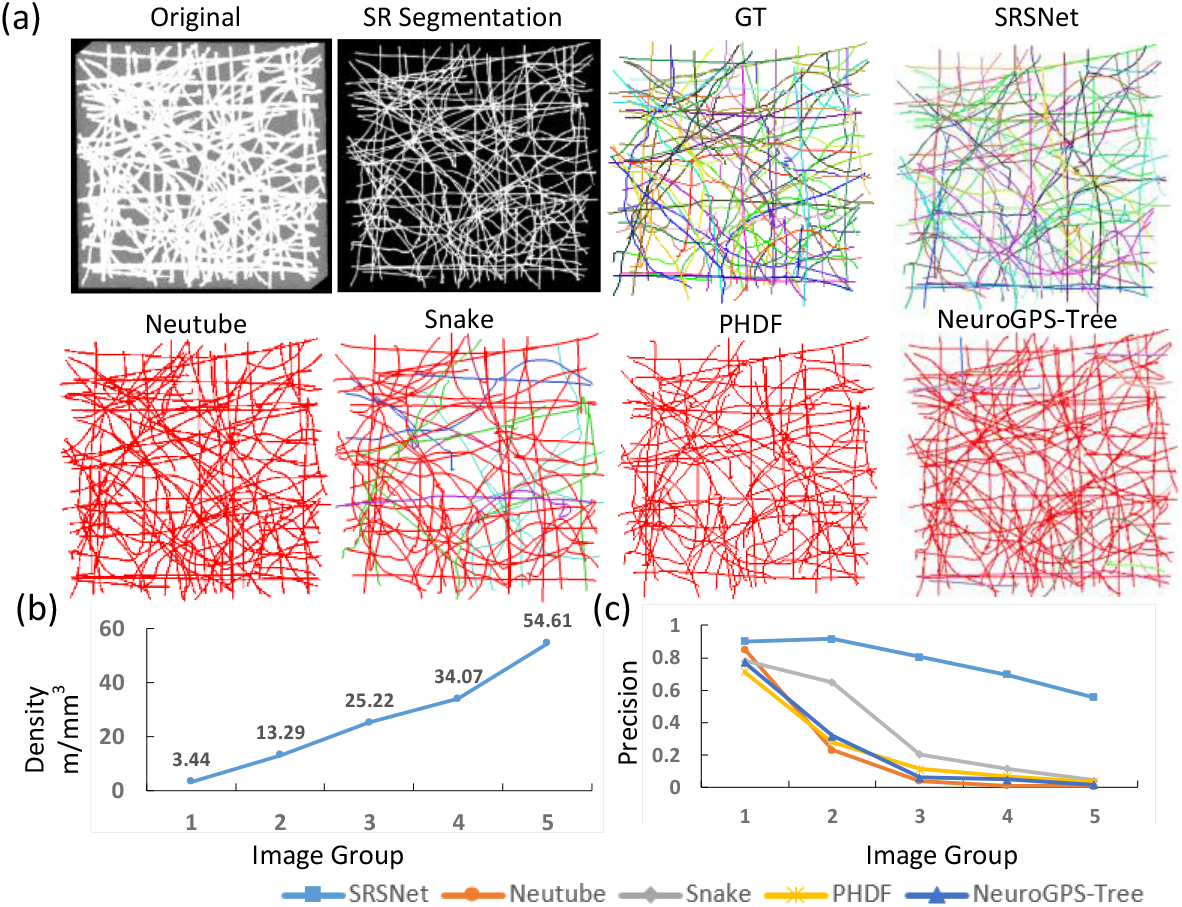
Super-resolution segmentation net (SRSNet) improves the reconstruction accuracy of packed neurites. (a) A synthesized chaotic image from group 3, the corresponding super-resolution image and the trace results obtained from super-resolution segmentation as well as that of four other automatic tracing methods from original image. The size of the simulation image is 256 × 256 × 128 voxels. (b) and (c) show the density and precision of neurites in five groups of synthesized chaotic images. The image groups are sorted according to the density of the neurites.

In **Fig. 2(c)**, all of methods achieve high recall value. It means that almost all of the skeleton of neurites are traced, although the other four methods cannot extract connection relationship correctly. In the case of the dense chaotic neurite image, these four tracing methods cannot be applied to tracing packed neurites.

We have evaluated the segmentation performance of SRSNet on mouse neuronal images. The testing images were acquired with fMOST [3]. The testing dataset includes 16 image stacks of dimensions 64 × 64 × 64 voxels. The resolution of neuronal images is 0.2 × 0.2 × 1 μm. The neurites in the image stacks are mainly axons and have a wide range of image intensities. The image stack and its segmentation results (**Figs. 3(a)**) derived from ResNet and SRSNet have been presented in **Fig. 3**. In ResNet, we have used the feature extracting network that contains four ResNet layers with 64 convolution kernel to replace the super-resolution head and the upsampling networks. The size of the convolution kernel is 3. This structure has been proposed for the voxel-wise residual network and has been widely applied in the segmentation of bio-images [20]. Compared to the segmentation done using ResNet, SRSNet maps the segmented neurites in a higher-resolution space and reduces the radius of the segmented neurites (**Figs. 3(b)**). We note that the segmented neurites driven from ResNet are thicker than in reality in most cases, which is partially due to the sharp changes in the image intensities. For quantifying the evaluation, the neurites have been manually traced and their traced skeleton points have been optimized. The skeleton points that were covered in the segmented regions generated with ResNet and SRSNet have been counted and the count rate has been plotted in **Fig. 3(c)**. The rate of the covered points was 0.96±0.02 for SRSNet and 0.99±0.01 for ResNet. Despite the existence of these slight differences, SRSNet still provides accurate segmentation. We have calculated the radii of the neurites along their manually traced skeletons in the original image stacks and their segmentation images. The calculated radii were sorted increasingly and normalized (**Fig. 3(d)**). From the results, we conclude that SRSNet can significantly reduce the effects of the point spread function and thus enlarge the distances between the boundaries of the neurites.

**Fig. 3.**
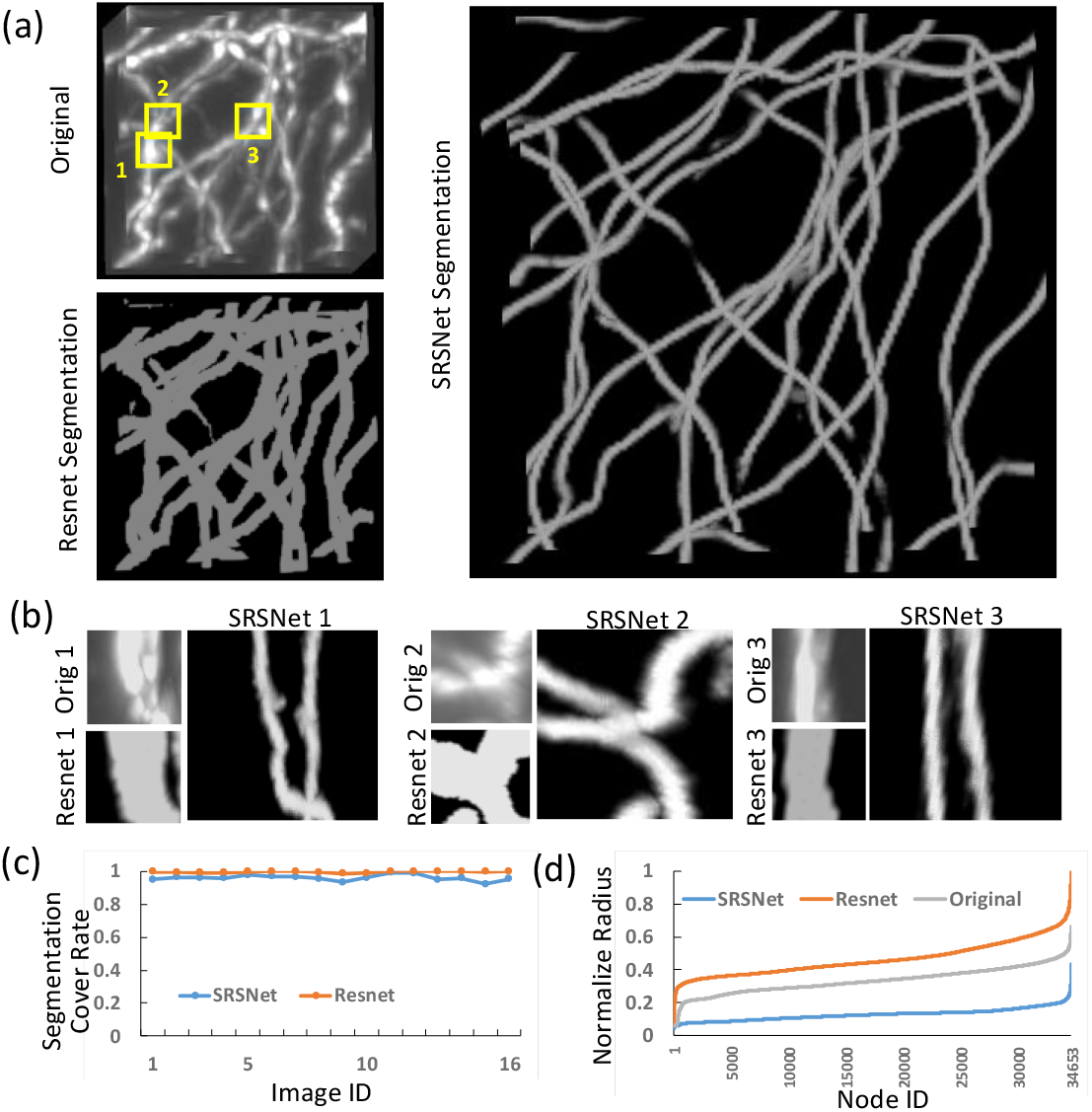
Comparison of the segmentation results obtained using SRSNet and ResNet-based method. (a) The neuronal image stacks has 64 × 64 × 64 voxels. (b) Segmentation obtained using a ResNet-based method which is widely applied in biomedical images. (c) Segmentation obtained using SRSNet. (d) The enlarged view of the subblocks and their segmentations indicated by yellow boxes 1-3 in (a). (e) The rates of the manually traced skeleton points of neurites dropping in the segmented region derived from SRSNet and ResNet-based method. (f) Results of the radii of the neuritis, with their centers at the manually traced skeleton points, calculated in the original image stacks, their segmentation image stacks generated by SRSNet and a ResNet-based method. The radii have been sorted and normalized.

Finally, we demonstrate that SRSNet essentially boosts the reconstruction accuracy of real packed neurites (**Fig. 4**). The testing dataset includes a stack of 16 images having dimensions of 128 × 128 × 64 voxels. One half is from the cortex region and the other half is from the hippocampus region. These image stacks include many densely distributed neurites. Their images intensities range from 7.36 m/mm^3^ to 27.03 m/mm^3^. The reconstruction tools such as Neutube, Snake, PHDF, and NeuroGPS-Tree have been used for illustrating the merits of SRSNet above these tools. Two image stacks have been shown and the reconstruction of neurites from them have also been presented in the figure. The results suggest that these tools can extract the skeletons of the neurites satisfactorily, but fail to build the links between the traced neurites. The false links sharply increase in a fixed space for these reconstruction tools as the number of neurites increase, as is indicated by arrows in **Fig. 1(a)**. In contrast, SRSNet significantly reduces the false links and generates accurate reconstructions in the case where the other reconstruction tools are powerless. We furthermore present the quantifying results. In the hippocampus datasets, the precision rate of SRSNet is 0.91±0.07, while it is 0.66±0.10 for NeuroGPS-Tree which provides the best reconstruction among these selected tools. SRSNet also attains a reasonable level of recall rate. On the other hand, both Neutube and PHDF tools generate the reconstruction with a high recall rate, however their precision in reconstruction is low. From these results, it can be understood that the skeleton of the traced neurites can be satisfactorily extracted and many traced neurites was falsely linked to each other, which is consistent with the two reconstruction examples. Similar results obtained for the reconstruction in the case of the cortex datasets (**Fig. 4(d) and 4(e)**) also suggest that SRSNet can achieve an accurate reconstruction of the packed neurites.

**Fig. 4.**
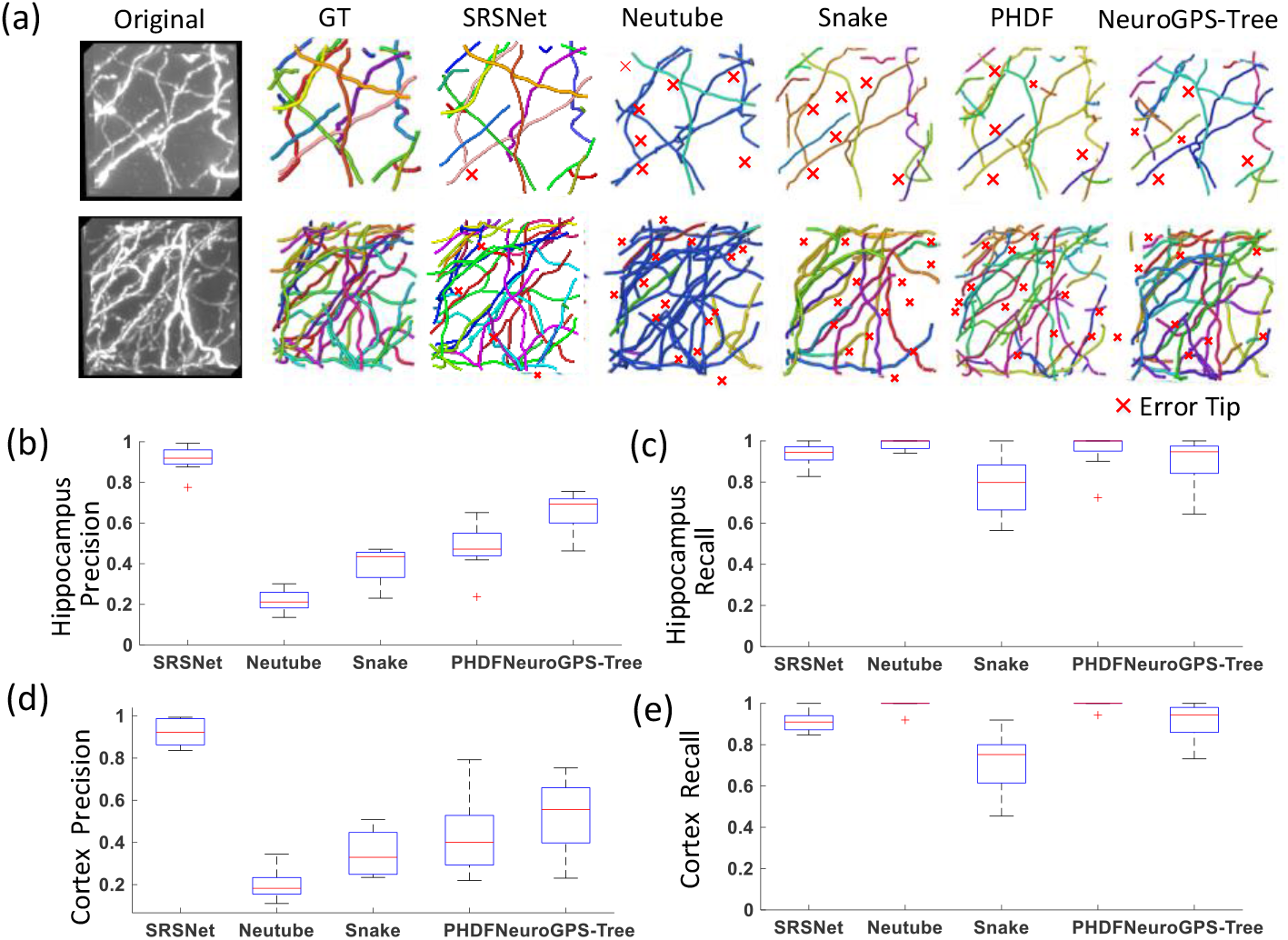
The reconstruction performance of SRSNet and four other automatic methods. (a) The two image stacks and their automatic reconstructions. The manual reconstruction is regarded as the ground truth (the second column). False links in the reconstruction are marked by red crosses. (b) and (c) show the precision and recall rates in the automatic reconstructions from 8 image stacks in the hippocampus region. (d) and (e) show the quantified reconstructions from 8 image stacks in the cortex region.

## Conclusions

In this work, we have identified that the small distance between the packed neurite boundaries and the crossover neurites that form various potential link patterns, are important factors which cause difficulty in tracing the packed neurites using the existing automatic tracing methods. We have proposed a 3D super-resolution segmentation network, called SRSNet, to acquire high-resolution segmentation images. The super-resolution segmentation images allow the automatic tracing methods to work in 3D HR space, resulting in high accuracy tracing. SRSNet can enlarge the 3D segmentation image by 16-fold. Simultaneously, SRSNet reduces the radii of the neurites by half owing to its super-resolution property, resulting in the enlargement of the distance between the boundaries of the neurites. In other words, it essentially improves the resolution of the neurites in an image. We have demonstrated the use of SRSNet in simulating cross neurites and parallel neurites and images of real dense neurites. The neighboring neurites are observed to be separated correctly in the super-resolution segmentation images. The precision of the tracing results on the super-resolution images has been found to be better than that of the automatic tracing methods applied on the original images. The 3D super-resolution segmentation method provides a new and effective way to solve the problem of dense neuron reconstruction.

